# Possible regulation of the immune modulator tetraspanin CD81 by alpha-synuclein in melanoma

**DOI:** 10.1101/2024.05.09.593218

**Authors:** Nirjhar M. Aloy, Christina Coughlan, Michael W. Graner, Stephan N. Witt

## Abstract

We probed the mechanism by which the Parkinson’s disease-associated protein α-synuclein (α-syn)/*SNCA* promotes the pathogenesis and progression of melanoma. We found that the human melanoma cell line SK-MEL-28 in which *SNCA* is knocked out (*SNCA*-KO) has low levels of tetraspanin CD81, which is a cell-surface protein that promotes invasion, migration, and immune suppression. Analyzing data from the Cancer Genome Atlas, we show that *SNCA* and *CD81* mRNA levels are positively correlated in melanoma; melanoma survival is inversely related to the levels of *SNCA* and *CD81*; and *SNCA*/*CD81* are inversely related to the expression of key cytokine genes (*IL12A*, *IL12B*, *IFN*, *IFNG*, *PRF1* and *GZMB*) for immune activation and immune cell-mediated killing of melanoma cells. We propose that high levels of α-syn and CD81 in melanoma and in immune cells drive invasion and migration and in parallel cause an immunosuppressive microenvironment; these contributing factors lead to aggressive melanomas.

**Graphical Abstract:** 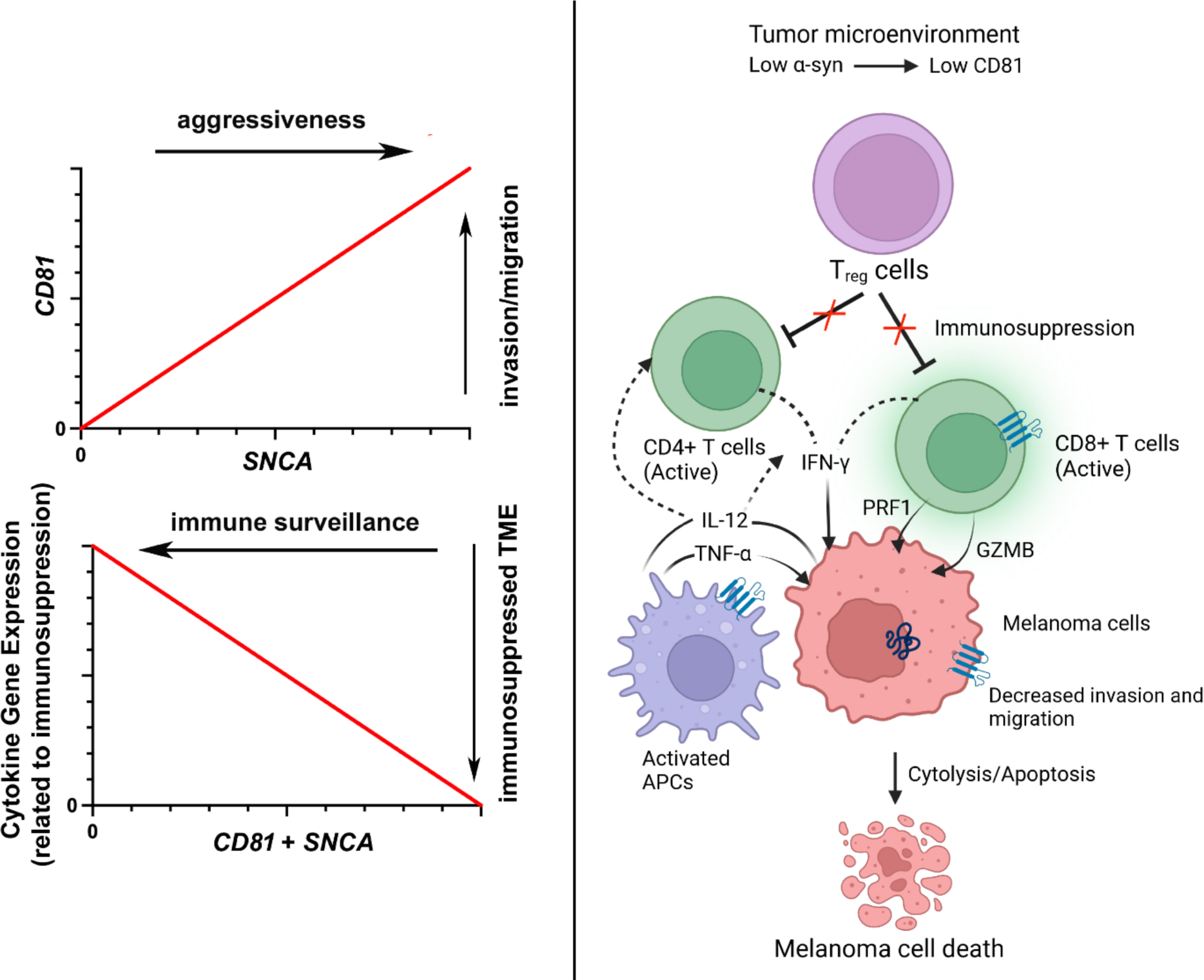

## 1. Introduction

Parkinson’s disease (PD) is a progressive neurodegenerative disease in which midbrain dopaminergic neurons die, leading to disturbances in gait and balance and the development of resting tremor [1]. Cancer is a disease of uncontrolled cell growth [2]. Although PD and cancer would seem to be unrelated, individuals with PD appear to be protected from all cancers except melanoma and brain cancer [3-5]. The co-occurrence of PD and melanoma is fueling a variety of studies to understand the molecular basis for this phenomenon. A recent meta-analysis of 63 publications, which included data obtained from over 17 million individuals, confirmed an inverse relation between PD and cancer, i.e., individuals with PD have a 15% lower risk of developing cancer, whereas individuals with cancer have a 26% lower risk of developing PD [6]. Strikingly, individuals with PD have a 75% higher risk of melanoma [6]. Several genes, including *SNCA*, may be involved in the co-occurrence of these two diseases [7,8]. Our focus here is *SNCA*, which codes for the presynaptic protein alpha-synuclein (α-syn) and which is highly expressed in neurons and melanocytes [9].

α-syn is an intrinsically unfolded protein [10] in solution that adopts an α-helical conformation when it binds to membranes enriched with negatively charged phospholipids (PL) [11,12]. Due to an internal hydrophobic domain, synuclein self-associates into a plethora of soluble oligomeric species, as well as insoluble amyloid fibers [13]. Some of these soluble amyloid fragments are likely the cytotoxic species that cause neurodegeneration in synucleinopathies like PD [14], dementia with Lewy bodies, and multiple system atrophy. In neurons, α-syn promotes SNARE complex assembly [15] and dilation of exocytic fusion pore [16].

In melanoma, α-syn functions in autophagy [17], melanin synthesis [18], and vesicular trafficking [19]. For example, α-syn inhibits the UVB light-induced increase in melanin synthesis, consequently melanin content is lower when melanoma cells express α-syn [18]. This implies that robust α-syn expression makes melanocytes more susceptible to UV-induced radiation damage. For unknown reasons, loss of α-syn expression in cutaneous melanoma cells results in decreased levels of cell surface proteins such as the transferrin receptor and the adhesion proteins L1CAM and N-cadherin [19].

Here we report that loss of α-syn expression in SK-MEL-28 cutaneous melanoma cells results in a significant decrease in the level of the cell surface protein CD81, and that α-syn and CD81 promote melanoma tumor progression by suppressing the immune response. CD81, which is a member of the tetraspanin family [20], promotes invasion and migration [21], immune suppression [22], and is a receptor for viral attachment [23]. Thus, alterations in α-syn and CD81 expression may have impacts on melanoma tumorigenicity at multiple levels.

## 2. Materials and methods

### 2.1. Cells and Cell Culture

SK-MEL-28 cells were purchased from ATCC (Catalog number HTB-72). *SNCA* was knocked out in our lab using CRISPR-Cas9 [24]. α-syn expression was rescued in KO8 and KO9 clones using lentivirus constructs, and the ‘knock-in’ clones were named KI8 and KI9, respectively. Cells were cultured and maintained at 37°C in Dulbecco’s Modified Eagle’s Medium (DMEM, ATCC-30-2002) that was supplemented with 10% fetal bovine serum (ATCC, 30-2020) and 1% penicillin-streptomycin.

### 2.2. RNA extraction and RT-qPCR

SK-MEL-28 cells were grown for 48 hours at 37°C, at 5% CO2 in complete DMEM medium. Total RNA was extracted using E.Z.N.A total RNA kit I (Omega Boi-Tek, cat.-R6834-02) following the manufacturer’s protocol. RNA quality and concentration was assessed with NanoDrop 2000c (Thermo Scientific). cDNA was synthesized from 1 μg of total RNA from each sample and qPCR and primer details are as described in the Supplementary Information.

### 2.3. Western Blotting

The preparation of cell lysates, LDS-PAGE, Western blotting, and densitometry were carried out as previously described [19,24]. Protein band intensities were measured using a BioRad Chemi-doc system, and intensities were normalized to the loading control β-actin. The detailed protocol is described in Supplementary Information.

### 2.4. Reagents and antibodies

We used the following mouse primary monoclonal antibodies: anti human CD81 (1:500, Santa Cruz Biotechnology), CD63 (1:500, Abcam), HSP70/HSP72 (1:1000, Enzo, C92F3A-5), α-syn (1:1000, Santa Cruz Biotechnology), α-Tubulin (1:4000, Sigma Aldrich). We used the following rabbit primary polyclonal antibodies: human CD9 (1:500, Systems Bioscience, # EXOAB-CD9A-1), HSP90 Antibody (1:1000, Cell Signaling Technology #4874). The secondary antibodies were Mouse–sc–516102 and Rabbit–sc– 2357 (1:2000, Santa Cruz Biotechnology).

### 2.5. Isolation of EVs

Melanoma cells were cultured in Dulbecco’s Modified Eagle’s Medium (DMEM, ATCC-30-2002) at 37°C that was supplemented with 10% exosome-depleted fetal bovine serum (Exo-FBS, Systems Bioscience) and 1% penicillin-streptomycin for 48 hours in T-75 flasks. 10 ml of conditioned media was collected from each flask and centrifuged at 3,000 g for 15 minutes to pellet live and dead cells, cell debris, and large microvesicles. The supernatant was collected for EV-isolation. 2 mL of ExoQuick TC ULTRA (Cat-EXOTC10A-1) reagent was added to the supernatant at 1:5 ratio and EVs were isolated using the manufacturer’s protocol.

### 2.6. Nanoparticle tracking

Purified EV samples were preserved at -80°C and sent to the University of Colorado Anschutz Campus for nanoparticle tracking particle analysis. The size distributions of EVs were determined by NTA Nanosight device (NS300, Software Version: NTA 3.2 Dev Build 3.2.16; Malvern Panalytical, Malvern WR14 1XZ, UK). Instrumental details are as described in Supplementary Information.

### 2.7 Bioinformatics

For the correlation and survival analyses of *CD81* and *SNCA* expression, we analyzed the skin cutaneous melanoma dataset from the TCGA PanCancer Atlas using cBioportal. The dataset contains the RNAseq data and the clinical features with outcomes. The Kaplan-Meier survival curve was constructed using the built-in survival analysis option of cBioportal [accessed on 2/25/24]. The high expression group was defined as the patient samples, in which the expression of both *SNCA* and *CD81* mRNAs were 0.5z score higher than the mean expression value. The expression values were in RSEM units (RNAseq by Expectation Minimization). Raw data may be obtained from cBioportal. Cytokine gene expression plots were created using GraphPad Prism v. 8 from the downloaded RNA-seq data from cBioportal [accessed on 3/20/2024].

### 2.8. Statistical Analysis

All the statistical analyses were performed in GraphPad Prism v. 8. Experiments were performed for 3 biological replicates. Figures were prepared in Adobe Illustrator v. 6.

## 3. Results

### 3.1. Loss of α-syn expression in SK-MEL-28 melanoma cells causes a significant decrease in the level of the cell-surface tetraspanin CD81

We previously found that knocking out *SNCA* causes reductions in motility, invasion, and migration compared to SK-MEL-28 melanoma control cells [19], and we attributed these effects to low levels of the adhesion protein L1CAM. Because CD81 is also involved in invasion and migration [19], we tested whether this protein is also downregulated in *SNCA*-KO clones. To this end, Western blotting revealed a significant reduction (*P* = 0.0188, *P* = 0.0109) in the level of CD81 in the lysates of *SNCA*-KO clones, KO8 and KO9, compared to control cells (Fig. 1A, B and Supplementary Fig. S1). The CD81 protein level significantly increased (*P* = 0.04) in KI9 cells compared to KO9 cells. The CD81 level increased in KI8 cells compared to KO8 cells, but the difference was not statistically significant (Fig. 1B).

**Fig. 1.**
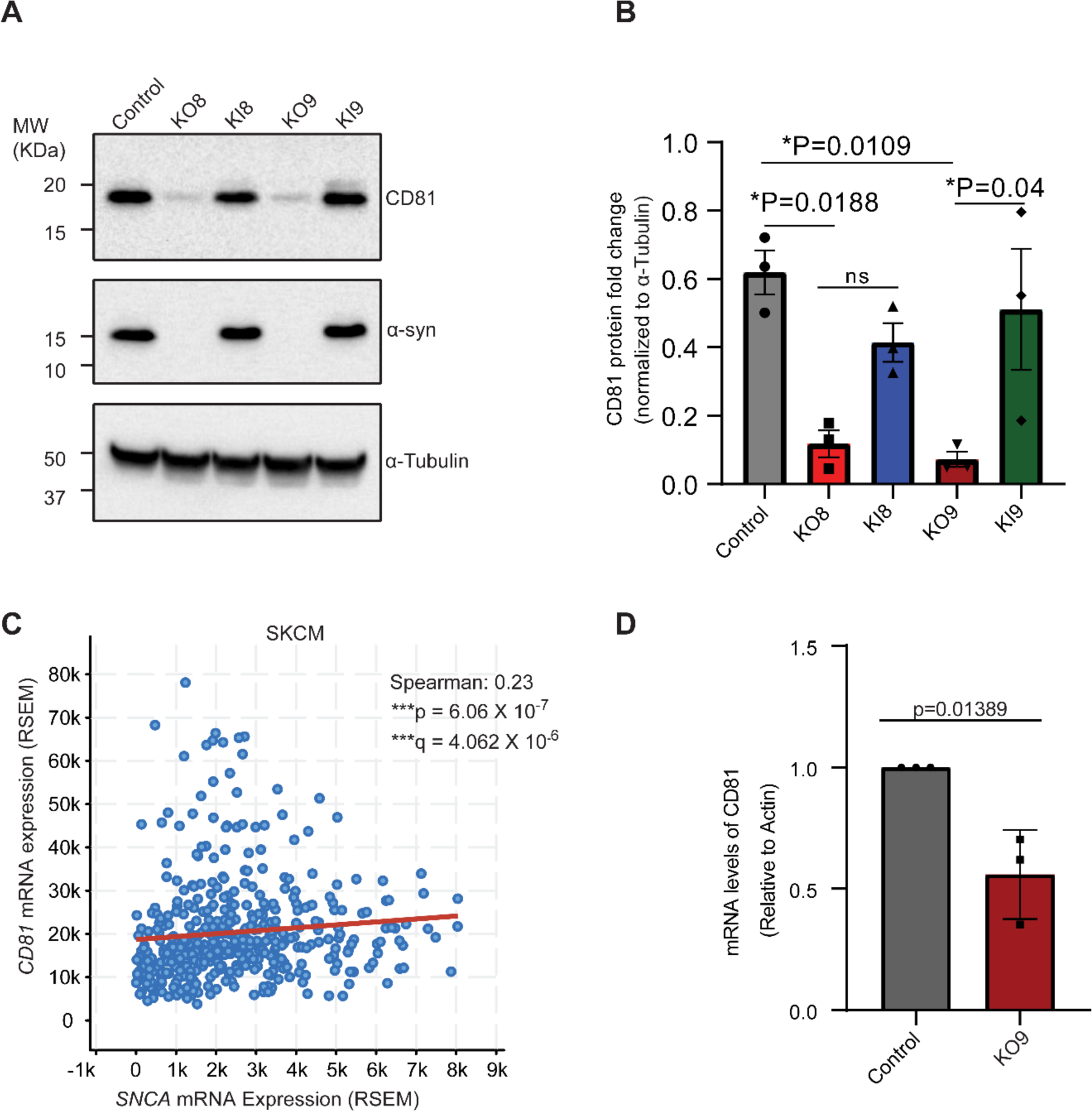
The level of the tetraspanin protein CD81 is decreased in *SNCA*-KO cells. (A) Representative Western blots of CD81 and α-syn in cell lysates. An equal amount of protein was loaded into each lane, followed by LDS-PAGE, and Western blotting. CD81 appeared as a band at 18 kDa, which is a mass consistent with isoform 2 (NP_001284578.1) [37]. (B) Quantification of band intensities. The band intensities were normalized to α-tubulin. Data are mean ± s.d. *P*-values were determined by one-way ANOVA with Tukey’s post hoc test (N=3). (C) Correlation of *SNCA* and *CD81* mRNA expression from RNAseq data in SKCM (Skin Cutaneous Melanoma dataset from the TCGA, PanCancer Atlas), N = 443 samples (samples with mRNA data). The correlation plots were generated by using the interactive platform of cBioportal and raw RNAseq data from the relevant datasets can be downloaded via cBioportal. Z-threshold=+/-0.1. RSEM=RNA-Seq by Expectation Maximization. (D) CD81 mRNA level is decreased in *SNCA*-KO melanoma cells. RT-qPCR showing the relative fold change of *CD81* mRNA level in SK-MEL-28 control and *SNCA*-KO cells. N = 3 biological replicates. Student’s paired t-test using GraphPad prism, version 8. Error bar, ± s.d.

We also conducted preliminary experiments on EVs released from control, KO, and KI melanoma lines. The EVs were characterized by Western blotting for common EV markers (CD81, CD9, CD63, HSP70, HSP90) and by nanoparticle tracking analysis (NTA) (Supplementary Fig. S2). We observed a decreasing trend in CD81 in *SNCA*-KO clone KO8 compared to control and KI clones (Supplementary Fig. S2). Future work will provide detailed quantitative information on the EV-cargos.

Based the above findings, we predicted that *CD81* and *SNCA* would be positively correlated in melanoma patient samples and perhaps in other cancers. To that end, analyzing RNAseq data in the TCGA databank, we found weak-to-moderate, and yet significant, positive correlations between *SNCA* and *CD81* in cutaneous melanoma (SKCM) (R = 0.23, P = 6.06 x 10^-7^) (Fig, 1 C) and in several other cancers (see Supplementary Fig. S3). We also conducted RT-qPCR analysis of SK-MEL-28 control and SK-MEL-28 *SNCA*-KO9 cells and found a significant decrease (41%, p = 0.0138) in *CD81* transcript level in KO9 cells compared to the control cells (Fig. 1 D). This latter result was consistent with our bioinformatics analysis of patient samples, which showed a direct relation (positive correlation) between *CD81* and *SNCA* mRNA levels.

### 3.2. *CD81* and survival

To address whether the expression of *SNCA* and *CD81* impacted the survival of melanoma patients, we analyzed the RNAseq data from melanoma patient samples (N = 443) from the skin cutaneous melanoma (SKCM) dataset of the PanCancer atlas [25]. We found that melanoma-specific survival was significantly lower in patients with elevated mRNA levels of *CD81* only (median survival 65.9 months versus 138.8 months, p = 0.0106, q = 0.0117) (Fig. 2 A). Interestingly, median survival was further decreased in patients with elevated mRNA level of both or at least one of *SNCA* and/or *CD81* (Fig. 2 B).

**Fig. 2.**
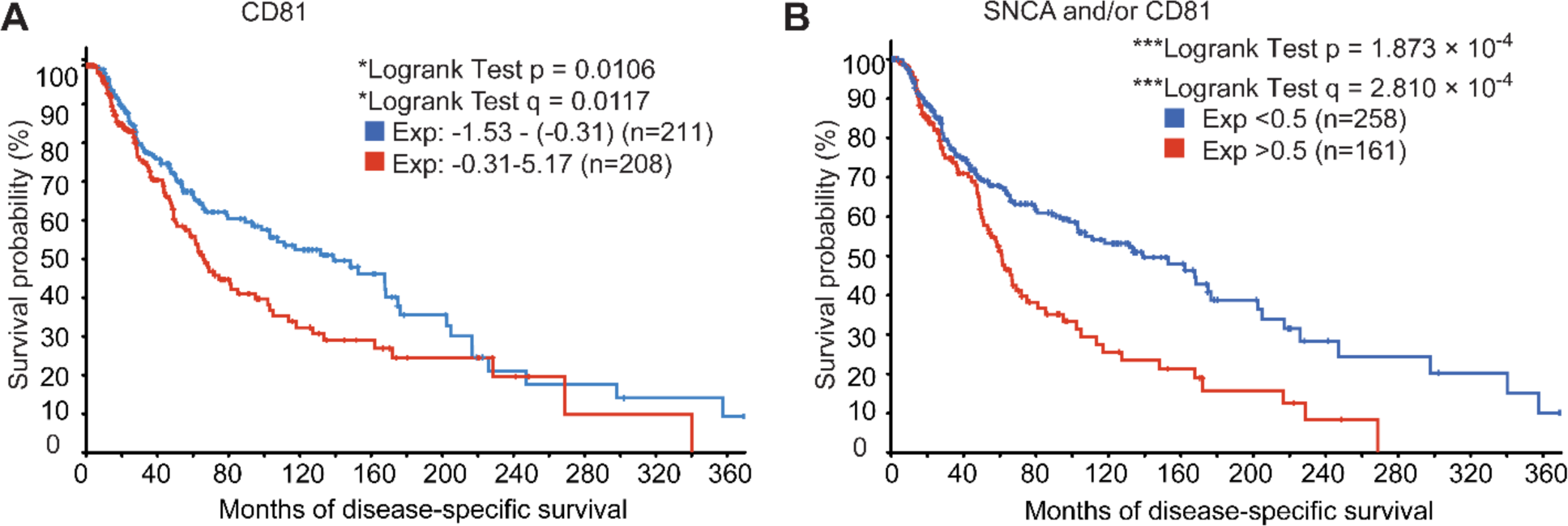
Melanoma-specific survival probability is decreased in patients with high expression of *SNCA* and *CD81* genes. (A) Melanoma-specific survival is decreased in patients with higher expression of *CD81*. Red line depicts melanoma patients with higher expression (above the median expression) of *CD81* (-0.31 - 5.17). Blue line depicts melanoma patients with lower expression (below the median expression) of CD81 (-1.53 - (-0.31). Median expression = -0.31. N = 419, Low expression group = 211, high expression group= 208. (B) Melanoma-specific survival is decreased in patients with higher expression of at least one or both of *SNCA* and *CD81*. The red line depicts melanoma patients with high expression (Exp > 0.5 SD above the median expression) of at least one or both of *SNCA* and *CD81*. (N = 161, both high group = 39, *SNCA* only high group = 70, *CD81* only high group = 52), and the blue line depicts patients with low expression (Exp < 0.5 SD below the median expression) of at least one or both of *SNCA* and *CD81* (n=258). Z threshold =+/- 0.5. The total number of patients in the analysis, N = 419. Disease-specific survival option was chosen from cBioportal to generate K-M survival curve.

### 3.3. CD81/SNCA and cytokines

α-syn and CD81 play important roles in the immune system. For example, (i) α-syn promotes neuroinflammation [26]. (ii) α-syn and CD81 are each involved in the developmental and functional modulations of B and T cells [27,28] (iii) A recent report showed immune cell mediated elimination of melanoma cells in the *SNCA*^-/-^ mouse model, which was associated with increased expression of *IFNG* and *TNF* [29]. This observation was consistent with a previous report on immune response from *SNCA*^-/-^ mouse showing increased production of IFN-γ in an experimental autoimmune encephalomyelitis (EAE) murine model [26]. (iv) Elevated expression of IFNG and TNF occurs upon blocking CD81 with antibody in mouse splenic cells [30]. Based on the above studies, we hypothesized that α-syn and CD81 play immunosuppressive roles in the tumor microenvironment (TME) of melanoma patients which leads to decreased survival.

To address whether decreased survival in patients with high expression of *SNCA* and *CD81* was associated with immunosuppression, we analyzed the mRNA expression of some cytokine genes related to immune activation (*IL12A*, *IL12B*, *IFNG, TNF*) [31,32] and immune cell-mediated killing of cancer cells (*GZMB*, *PRF1*) [33]. In the same cohort of patients with decreased survival as described above (N = 174, 13 of these patients were not included in the survival study in Fig. 2B), we found significantly lower mRNA expression levels of *IL12A*, *IL12B*, *IFNG*, *TNF*, *PRF1*, and *GZMB* (Fig. 3 A-F). Taken together, the analysis in Fig. 3 reveals an intriguing inverse relation (negative correlation) between *SNCA*/*CD81* and cytokine genes.

**Fig. 3.**
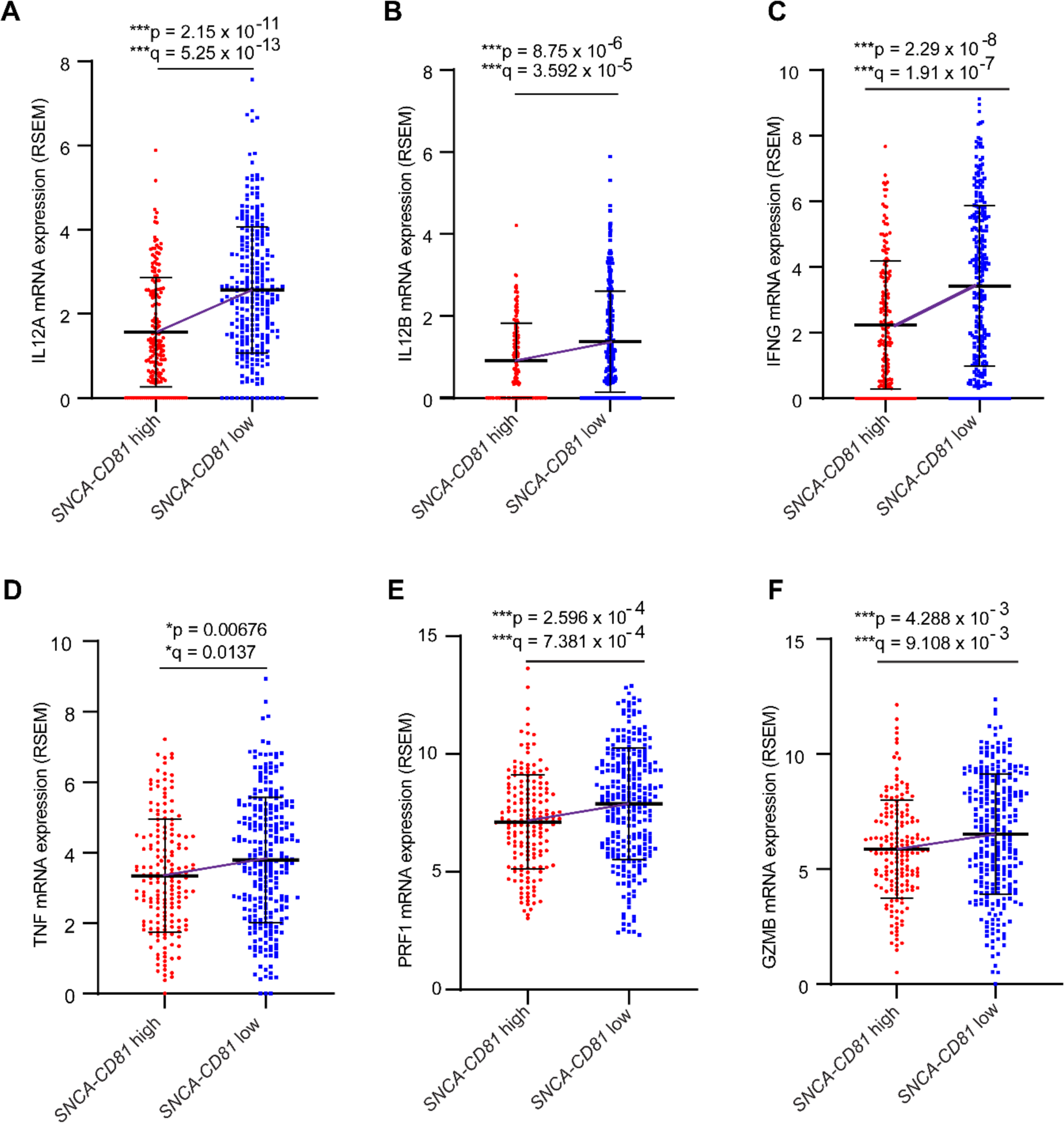
Elevated expression of cytokine genes in melanoma patient samples with low expression of *SNCA* and *CD81*. Patient groups: *SNCA-CD81* high: melanoma patients with high expression (Exp > 0.5 SD above than the median expression) of at least one or both of *SNCA* and *CD81* (n= 174, *SNCA* high 109, *CD81* high group = 91, both high = 26). *SNCA-CD81* low: melanoma patients with low expression (Exp < 0.5 SD below the median expression) of at least one or both of *SNCA* and *CD81* (N = 269)). Panels (B-F) focus on the same cohort of patients (N = 174) of which 13 patients were not included in the survival study by the study conductors. q = Benjamini-Hochberg estimate of false discovery rate. Red lines show the increase in the median values between groups.

## 4. Discussion

In this work we found that the loss of α-syn expression in SK-MEL-28 melanoma cell line results in a significant decrease in the level of the tetraspanin CD81 (Fig. 1). We previously discovered that loss of α-syn expression in this cell line downregulates the levels of several cell-surface proteins, i.e., TfR1, L1CAM, FPN1 (ferroportin), N-Cadherin [19,24]. We hypothesize that loss of α-syn expression causes the altered trafficking of some of these membrane proteins to the lysosome [19,24], where they are degraded. We have observed a 40% decrease in the mRNA abundance of CD81 and an 83% decrease in CD81 protein levels in each KO clone. This suggests that α-syn might have a role in both the regulation of the mRNA level and the trafficking of CD81. So, it is possible that α-syn is involved in modulation of CD81 at transcriptional, post-transcriptional and post-translational levels.

SK-MEL-28 *SNCA*-KO cells have significant defects in single-cell motility, invasion, and migration relative to control cells that express synuclein [19]. We reasonably concluded that these defects are a consequence of the *SNCA*-KO cells having a low level of L1CAM, which is a protein implicated in signaling, motility, and invasion [34]. Given that *SNCA*-KO cells also have a significantly low level of CD81 relative to control cells, and that CD81 promotes the invasiveness of melanoma cells, we conclude that the decrease in the motility, invasion, and migration of *SNCA*-KO cells is a consequence of low levels of both CD81 and L1CAM.

An intriguing finding is that high levels of both *SNCA* and *CD81* mRNA correlate with an immunosuppressive mRNA signature (Figs. 3, 4). Our results show an inverse relation between *SNCA*/*CD81* and cytokine genes, i.e., patients with high expression of both *SNCA* and *CD81* have decreased expression of key cytokine genes and decreased survival. Whereas patients with low expression of both *SNCA* and *CD81* have increased expression of key cytokine genes and increased survival. Based on our findings and those from previous studies, we developed a model to explain how *SNCA* and *CD81* influence survival (Fig. 4). Several points about this model are as follows: (i) Using a mouse model of breast cancer, S. Levy and colleagues by [22] showed that CD81 promotes tumor growth and metastasis by supporting the suppressive function of regulatory T (T_reg_) cells and myeloid-derived suppressor cells (MDSC). The same group showed that *CD81* was not necessary for Treg development but was necessary for the suppressive function of T_reg_ cells and promotes tumor growth. Loss of CD81 expression in our model disrupts T_regs_ and consequently activates CD4+ and CD8+ T cells and leads to increased expression of perforin (*PRF1*) and granzyme B (*GZMB*) and the eventual killing of melanoma cells. (ii) *IFNG* and *TNF* expression were increased in immune cells from *SNCA*^-/-^ mice [26,29] and in murine splenic cells upon treatment with antibody against CD81 [30]. (iii) Extracellular vesicles released from melanoma cells might be the key to maintaining or disrupting the suppressive function of T_reg_ and MDSC cells. If melanoma-derived EVs, which contain large numbers of CD81 and α-syn molecules with their differential cargos, are taken up by T_reg_ and MDSC cells, then the suppressive function of these immune cells could be strengthened and/or prolonged. Conversely, if melanoma-derived EVs, which contain low numbers of CD81 and α-syn molecules with their differential cargos, are taken up by T_reg_ and MDSC cells, then the suppressive function of these immune cells could be disrupted [35,36].

**Fig. 4.**
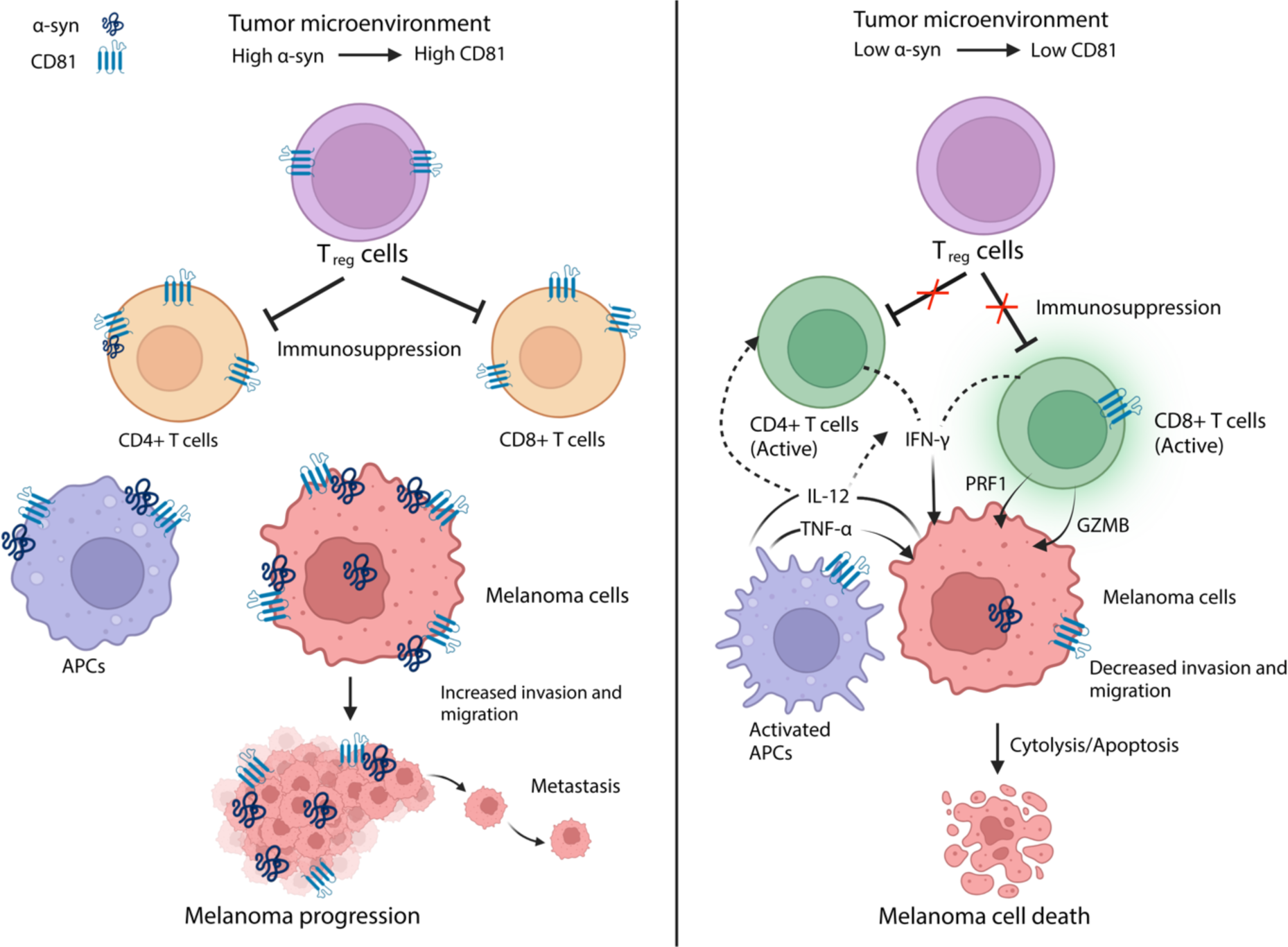
Proposed model for SNCA and CD81 modulating the TME. Left panel: elevated levels of SNCA and CD81 create an immune-suppressive TME. Right panel: Low levels of SNCA and CD81 activate Treg, CD4+, and CD8+ cells, resulting in cell-mediated killing of tumor cells. Created with BioRender.com

In conclusion, adding to the mutational burden, α-syn and CD81 promote the pathogenesis and progression of melanoma in two ways: (i) a high level of α-syn promotes the trafficking of cell surface proteins, such as L1CAM, TfR1, and CD81, to the cell surface, which promotes invasion and migration. (ii) high levels both α-syn and CD81 appear to inhibit the production of cytokines that are vital for immune activation and surveillance, which results in melanomas that essentially evade the immune system.

## Abbreviations

CRISPR: clustered regularly interspaced short palindromic repeats
GZMB: granzyme B
IL: interleukin
IFNG: interferon gamma
NK: natural killer cells
PD: Parkinson’s disease
PRF1: perforin 1
SKCM: cutaneous melanoma
TCGA: the cancer genome atlas program
TME: tumor microenvironment
TNF: tumor necrosis factor alpha

## Acknowledgements

We thank the Feist-Weiller Cancer Center for providing seed funds to SNW for his melanoma work and for a Carroll Feist Predoctoral Fellowship award (149741440A) to NMA. We thank Dr. Andrew D. Yurochko for the gift of the actin primers and the staff of the LSUHS Research Core Facility for their help with the instruments used in this work.

## Author contributions

Conceptualization: N.M.A., and S.N.W.; Investigation and data analysis: N.M.A., C.C., M.W.G., and S.N.W.; Project management: S.N.W.; Writing – original draft: N.M.A, and S.N.W.; Writing – review and editing: N.M.A., C.C., M.W.G., and S.N.W.

## Notes

### Competing Interest Statement

The authors have declared no competing interest.

### Summary of Updates

Figure 3 revised shows an inverse relation between SNCA/CD81 and cytokine genes IL12A, IL12B, IFNG, TNF, PRF1, and GZMB. This is an update of Figure 4C-F original. The CXCL13 correlation plot (Fig. 4D, original), which is correct, was not included in the revised version. Figure 4 revised is our model for how SNCA and CD81 affect melanoma cells and the tumor microenvironment. This figure was not in the original. Figure 3 original was deleted in the revised.

