## Supplemental info and figures for "Possible regulation of the immune modulator tetraspanin CD81 by alpha-synuclein in melanoma"

**for**

**Materials and Methods**

*Western Blotting*

SK-MEL-28 control, KO8, K09, KI8, KI9 cells were cultured for 48 hours in Dulbecco's Modified Eagle's Medium (DMEM, ATCC-30-2002) supplemented with 10% exosome depleted fetal bovine serum (Exo-FBS, Systems Bioscience) and 1% penicillin-streptomycin. After two washes with ice-cold PBS, cells were lysed in ice-cold RIPA buffer, and after passing through a 25G needle, the cell lysates were centrifuged at 18,360 g at 4°C for 20 minutes. Protein concentration was determined using the BCA protein assay (Thermofisher, Cat- 23250). Equal amounts of protein were loaded into each lane of NuPAGE^TM^ 4-12%, Bis-Tris mini protein gels (ThermoFisher, Cat-NP0336PK2) and PAGE was performed at constant voltage (170 watts). Semi dry transfer to nitrocellulose blots was performed using Trans blot Turbo (Biorad, Cat 1704150). After one hour of blocking in membrane blocking agent (G-Biosciences, Cat- 786-011), incubation with primary antibodies was done overnight at 4°C. After washing the membranes 5 times (5 minutes each) with PBST, incubation with secondary antibody was performed at room temperature. Western blot images were acquired using chemiDoc Touch machine (Biorad). The protein concentration of EV samples was determined using the BCA protein assay. EV samples were lysed with NuPAGE LDS sample buffer (Invitrogen, Cat – NP0007), and 20 μg of EV-protein was loaded into each lane of the polyacrylamide gel, and Western blotting was performed as described above. In some cases, membranes were stripped (G Biosciences, Cat- 786-305), cut into pieces, and the pieces were re-probed.

*RNA extraction and RT-qPCR*

SK-MEL-28 cells were grown for 48 hours at 37°C, at 5% CO2 in complete DMEM medium. Total RNA was extracted using E.Z.N.A total RNA kit I (Omega Boi-Tek, cat.- R6834-02) following the manufacturer’s protocol. RNA quality and concentration was assessed with NanoDrop 2000c (Thermo Scientific). cDNA was synthesized from 1 μg of total RNA from each sample as follows: 1 μg total RNA + 2 μL 5X first strand buffer + 1 μL DNAse I + DEPC treated ddH2O to make 10 μL total volume. This was incubated for 30 minutes at 37°C. Then, 1 μL of DNAseI-DNAse I-inhibitor was added and incubated at 65°C for 10 minutes and quenched on ice for 1 minute. The reverse transcription reaction was set up as follows: 5 μL DNAase treated RNA+1 μL 5X RT buffer + 0.25 μL random primers (1:30 dilution) + 0.25 μL dNTPs + 0.5 μL DTT + 0.2 μL RNAse-inhibitor + 0.2 μL MMLV RTase (Promega, cat. no. M1708) or (ddH20 for RT-) + 2.6 μL DEPC treated ddH2O = 10 μL final volume of mastermix was incubated at 37°C for 1 hour. This reaction was incubated at RT with a no primer control for each sample to ensure that there was no non-specific amplification in the qPCR reaction. Finally, the samples were treated with RNAseH at 37°C for 30 minutes.

qPCR was performed using the iTaq Universal SYBR Green Supermix (Bio-Rad, Cat. #1725121) according to the manufacturer’s protocol. Primers (Int. DNA Tech.):

CD81 Forward: 5’-CAAGTACCTGCTCTTCGTCTTC-3’

CD81 Reverse- 5’- TTGTCTCCCAGCTCCAGATA-3’

Actin: Forward: 5'-GGAGAAGAGCTACGAGCTGCCTGAC-3'

Actin Reverse: 5'-AAGGTAGTTTCGTGGATGCCACAGG-3'

*Nanoparticle tracking*

The EV-samples were preserved in Buffer B (Systems Bioscience) at -80°C and sent to the University of Colorado Anschutz Campus for nanotracking particle analysis (NTA). The size distributions of EVs were determined by NTA Nanosight device (NS300, Software Version: NTA 3.2 Dev Build 3.2.16; Malvern Panalytical, Malvern WR14 1XZ, UK). One ml of exosome sample was prepared by freshly diluting the exosome sample 1:1000; and 1:10,000 in ddH2O (0.1μm filtered) before reading. Exosome number was captured using the following analytical settings on the NTA: camera type: sCMOS, level 5 (NTA 3.0 levels); green laser; slide setting and gain: 600, 300; shutter: 15 ms; histogram lower limit: 0; upper limit: 8235; frame rate: 24.9825 fps; syringe pump speed: 25 arbitrary units; detection threshold: 7; max jump mode: auto; max jump distance: 15.0482; blur and minimum track length: auto; first frame: 0; total frames analyzed: 749; temp: 21.099-22.015 °C; viscosity: 1.05 cP.

**Figure S1**

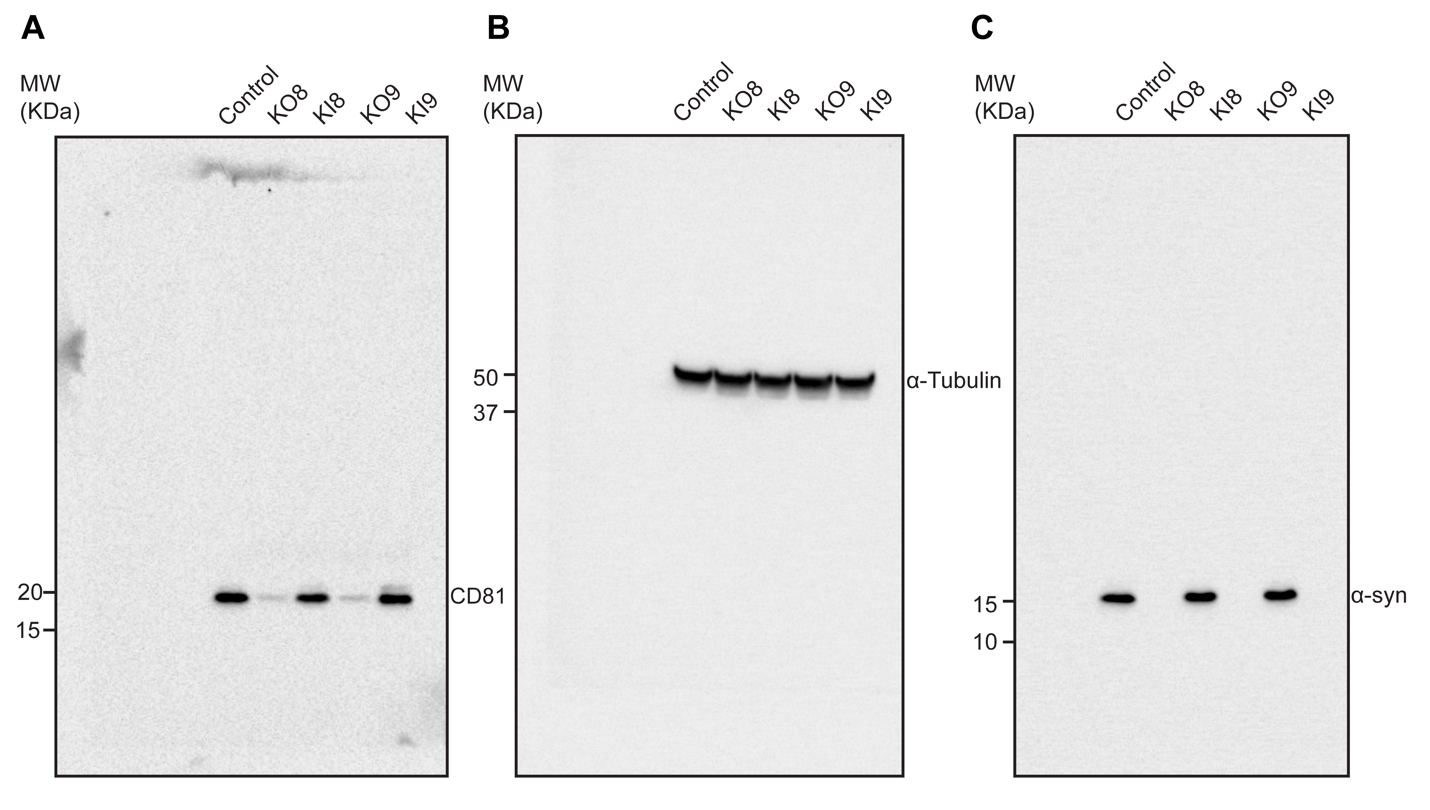

**Fig. S1** (A-C) Full-length original uncontrasted Western blots are shown cropped in Fig 1A.

**Fig. S2**

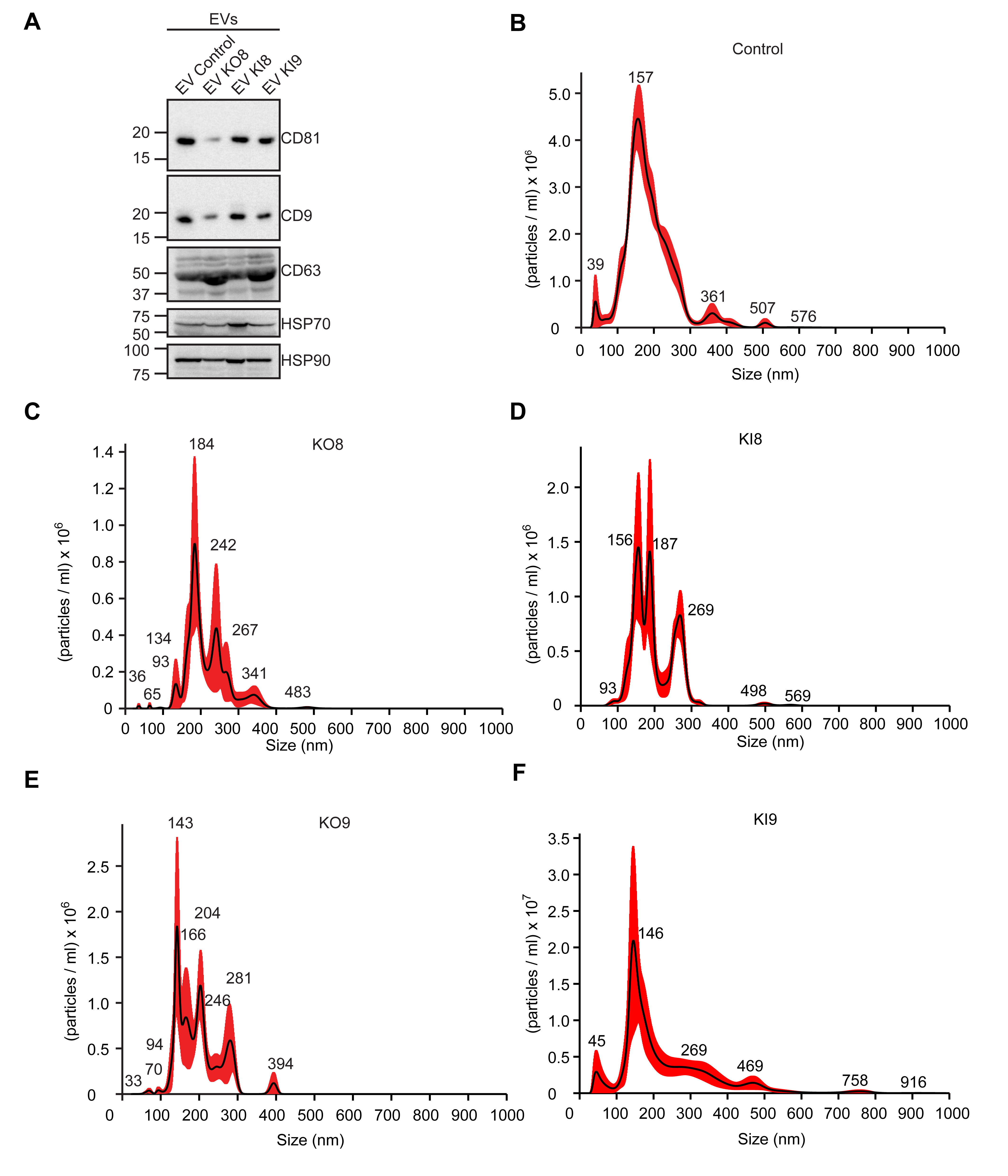

**Fig. S2** (A) Characterization of EVs by western blotting for common EV-markers. (B-F) NTA showing size-distribution of EVs isolated from SK-MEL-28 (B) Control, (C) KO8, (D) KI8, (E) KO9 and (F) KI9 cells. NTA images are representative of at least 3 independent experiments. Experiments were described in method section.

**Fig. S3**

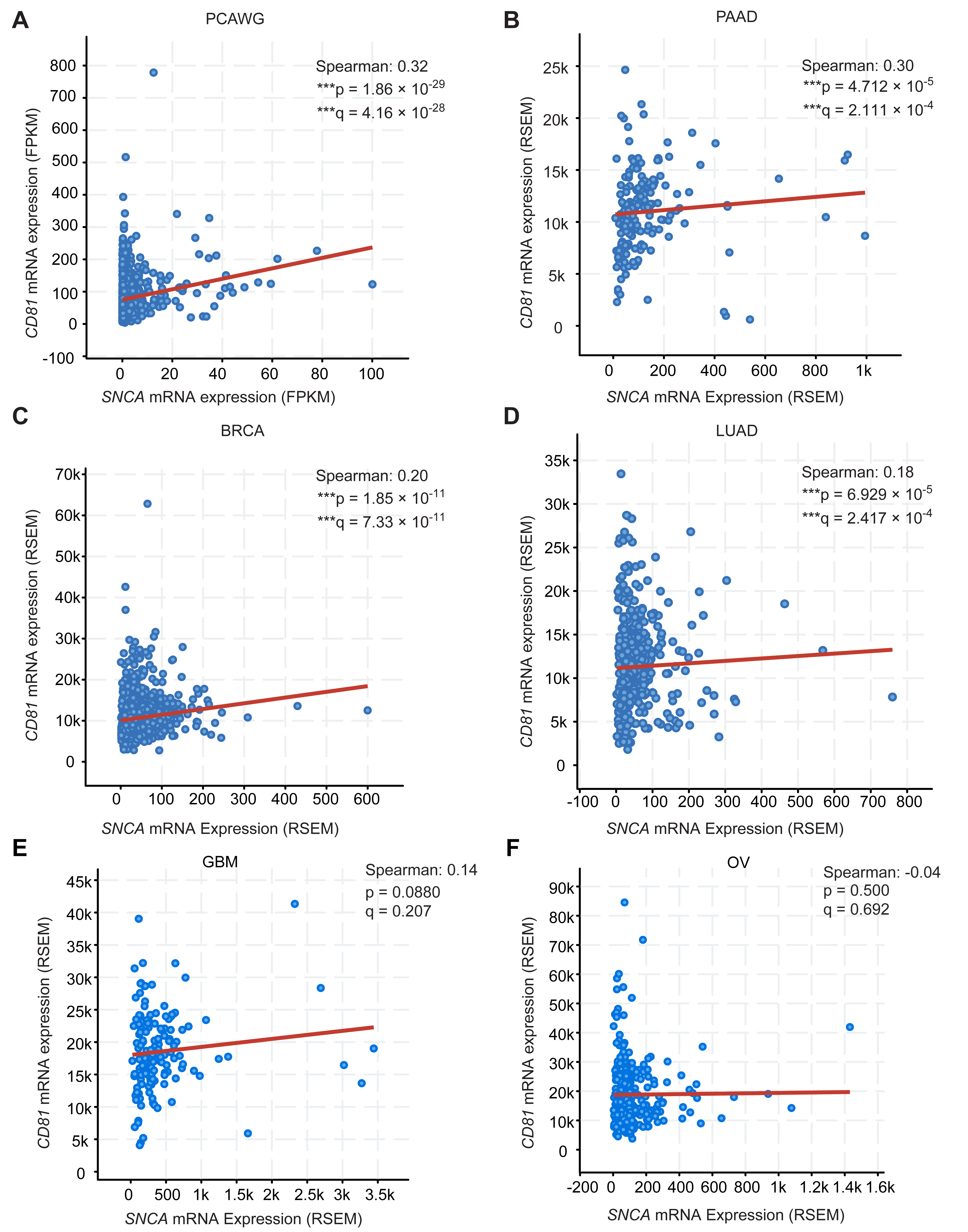

**Fig. S3** Correlation of mRNA expression levels of *SNCA* and *CD81* in different cancers. Bioinformatic analysis of RNAseq datasets suggests a weak correlation between the mRNA levels of *SNCA* and *CD81* in (A) PRAD (Prostate Adenocarcinoma) dataset. N = 493. (B) HCC (Hepatocellular Carcinoma) dataset, N = 366. (C) mRNA expression of *SNCA* and *CD81* have a non-significant correlation in GBM (Glioblastoma Multiforme) dataset, N = 160. (D) mRNA expression of *SNCA* and CD81 have no correlation in OV (Ovarian Serous Cystadenocarcinoma) dataset, N = 300.
